# Evo-engineering and the Cellular and Molecular Origins of the Vertebrate Spinal Cord

**DOI:** 10.1101/068882

**Authors:** Ben Steventon, Alfonso Martinez Arias

## Abstract

The formation of the spinal cord during early embryonic development in vertebrate embryos is a continuous process that begins at gastrulation and continues through to the completion of somitogenesis. Despite the conserved usage of patterning mechanisms and gene regulatory networks that act to generate specify spinal cord progenitors, there now exists two seemingly disparate models to account for their action. In the first, a posterior localized signalling source transforms previously anterior-specified neural plate into the spinal cord. In the second, a population of bipotent stem cells undergo continuous self-renewal and differentiation to progressively lay down the spinal cord and axial mesoderm by posterior growth. Whether this represents fundamental differences between the experimental model organisms utilised in the generation of these models remains to be addressed. Here we review lineage studies across four key vertebrate models: mouse, chicken, *Xenopus* and zebrafish and relate this to the underlying gene regulatory networks that are known to be required for spinal cord formation. We propose that by applying a dynamical systems approach to understanding how distinct neural and mesodermal fates arise from a bipotent progenitor pool, it is possible to begin to understand how differences in the dynamical cell behaviours such as proliferation rates and cell movements can map onto conserved regulatory networks to generate diversity in the timing of tissue generation and patterning during development.

## Introduction

The chordate central nervous system has a stereotypical organization along the anteroposterior axis. At the anterior end, a large mass of neuronal circuits, the brain, processes information from the rest of the body and determines and controls motor, sensory and cognitive functions. Behind the brain and extending posteriorly the length of the organism is the spinal cord that relays information to and from the brain to the rest of the body. The brain and the spinal cord are functionally and structurally continuous and form a most important physiological unit. Understanding their origin will provide insights into the emergence of circuits and their relationship to the rest of the organs in the body.

Much of our appreciation of the development of the nervous system is derived from interpretations of two experiments in amphibian embryos. The first one is the famed Spemann-Hilde Mangold manipulations of the organizer, a multicellular arrangement that appears at the beginning of gastrulation which has the property of evoking neural tissue on adjacent undifferentiated ectoderm (Spemann, 1924); these experiments led to the notion of neural induction. The second one was performed by P. Nieuwkoop, involved setting up interactions between different tissues around gastrulation and led to the ‘activation-transformation model’ (Nieuwkoop, 1954). According to this, the organiser induces neural tissue with anterior characteristics in the overlying ectoderm (*activation*) and a second polarized signal creates a gradient along the AP axis that promotes the posteriorisation (*transformation*) of the neural plate in a graded manner. This mechanism has been extended to all vertebrate embryos with discussions focusing more on the molecules mediating each of the steps than on the events themselves. However, a cellular perspective of the process reveals a number of issues, particularly of whether the activation/transformation model applies to all vertebrate embryos (Stern et al., 2006). Significantly inspection of the fate map of different embryos reveals a problem (Fig. 1). In all species, the brain is represented proportionately to the rest of the organism but there are significant differences in the fate map of the spinal cord. Whilst in anamniotes (e.g frogs and fish) the gastrula ectoderm contains a region of cells already fated to do this that is proportional to its target tissue, in amniotes (e.g chick and mouse) this region is small relative to the rest of the body. While it is easy to see how in anamniotes a transformative agent could act early on to specify different fates in a preexisting spinal cord, this cannot occur in amniotes. Simply: the spinal cord is not represented in the amniote gastrula in a proportionate manner to the rest of the body. Therefore, amniotes must make use of a different strategy to generate the spinal cord which must involve massive, but controlled, expansion of a progenitor population.

**Figure 1:**
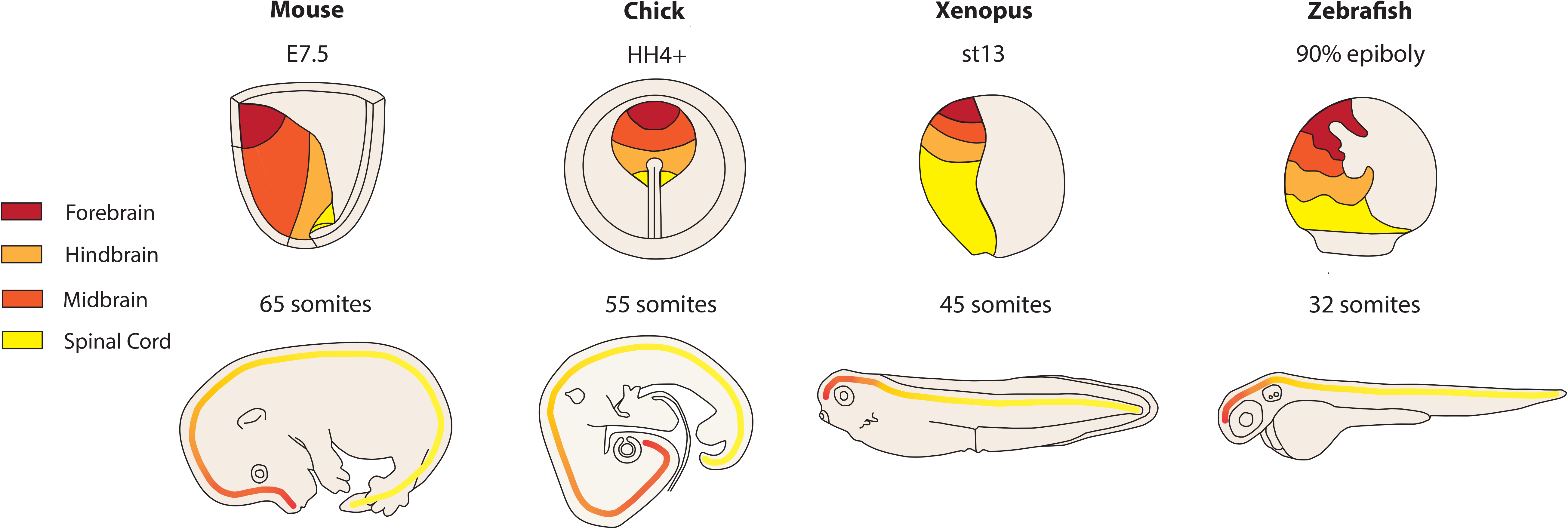
Fate maps of the prospective spinal cord region in four key vertebrate experimental models: mice, chick, *Xenopus* and zebrafish. On the upper panel, representative fate maps are shown for each species at roughly equivalent stages that mark the end of primary gastrulation. The entire region fated to generate the central nervous system is shown, colour coded according to prospective anterior-posterior level. Below are these regions mapped onto embryos after the completion of somitogenesis.

A difference between amniotes and anamniotes is emphasized by cell lineage tracing experiments in chick (Brown and Storey, 2000; Selleck and Stern, 1991) and mouse (Cambray and Wilson, 2007, 2002; Mathis and Nicolas, 2000; McGrew et al., 2008; Tzouanacou et al., 2009)) that have revealed a population of cells with stem/progenitor characteristics that generates the spinal cord directly, with little dual contribution to the anterior and posterior aspects of the central nervous system. More recently, a large-scale retrospective lineage analysis study in mouse has confirmed that, in mice, the spinal cord is derived from a stem/progenitor cell population and revealed that individual cells in this population can contribute to both spinal cord and paraxial mesoderm during axial extension (Tzouanacou et al., 2009). It has been suggested that this progenitor population can be identified by a co-expression of Sox2 and Brachyury/T and has been called “neuromesodermal progenitors” (NMps; Henrique, Abranches, Verrier, & Storey, 2015; Wilson, Olivera-Martinez, & Storey, 2009). Whether a similar population with self-renewal potential exists in anamniotes and how it relates to the anterior neural ectoderm is open to discussion (Martin and Kimelman, 2012; Steventon et al., 2016). Notwithstanding these considerations, the observations suggest that the brain and the spinal cord have different and maybe independent embryological origins in vertebrates.

Over the last few years there has been some progress in understanding the origin of the spinal cord and here we focus on this subject and recent developments. Comparing amniotes and anamniotes we address the question of whether the development of their nervous systems follow similar paths and suggest that in anamniotes the spinal cord is derived from a NMp population which, in contrast to that of amniotes, does not expand. Despite this difference, the molecular mechanisms required for the posterior extension of the embryo appear to be well conserved across chordates and to some degree across metazoans (Martin and Kimelman, 2009). In order to attempt to reconcile these seemingly disparate views on the degree of conservation in the mechanisms underlying spinal cord development, we go on to briefly review recent data analyzing the role of signal transduction networks in the maintenance and differentiation of NMps. Finally, we propose that by considering NMps as a transition state and modelling their development as a dynamic system, it may be possible to understand how evolution has acted up the signal and gene regulatory networks in order generate diverse cellular outputs from seemingly conserved molecular mechanisms.

## Neuromesodermal progenitors in amniotes: a stem cell pool

Neuromesodermal progenitors are often defined on the basis of co-expression of T/Bra, Sox2 and Nkx1.2 (Henrique et al., 2015). However, gene expression patterns are markers for cells rather than real identifiers. These, in the context of stem/progenitor cells should rely on the demonstration of stem/cell progenitor population clonogenicity. Thus, in order to assess how progenitor pools expand to generate the spinal cord, we must be able to label single cells and follow their derivatives through the entirety of embryonic development. In the case of mouse development, this has been achieved by retrospective clonal analysis using a LacZ transgene bearing an internal duplication that creates a frameshift (LaacZ) and inactivates the ß-galactosidase enzyme, for genetic lineage tracing. The LaacZ gene is placed under the control of a ubiquitous CNS promoter and rare spontaneous deletions recover the frame and the activity in single cells, generating long lived clones that thus reveal the modes of growth of progenitor pools during the development of the CNS (Mathis and Nicolas, 2000). These experiments produced regionalised clones, backdated to around E6.5 onwards, with distinct modes of growth. Long clones are restricted to the spinal cord with frequencies of clone lengths distributed in such a manner that suggest a clonal mode of growth. Anterior CNS clones appeared as orderly intermingled, in which cells can only rearrange with their closest neighbours (Mathis and Nicolas, 2000). The clonal growth of the spinal cord was paralleled by observations in which LaacZ expression with the acetylcholine receptor revealed a similar clone organization in the developing myotome (Nicolas et al., 1996). In similar experiments using the ubiquitous ROSA26 promoter, Tzouanacou and colleagues (Tzouanacou et al., 2009) provided evidence for a bipotent progenitor pool that gives rise both the spinal cord and paraxial mesoderm derivatives which also follow clonal distributions that are indicative of a stem mode of growth. In all cases the clones are anchored on the tail region of the embryo and suggest the existence of a stem/progenitor population in this region. These experiment provided evidence for the existence of NMps and for the way their activity is coordinated over time.

A key feature of a *bona fide* stem cell population is the ability for long term self-renewal which contrasts with the temporally limited expansion of the NMp pool during embryogenesis.

However, transplants of cells from the posterior region of older embryos into younger host demonstrated that cells within this region retain the ability to generate progenitors of both the spinal cord and mesoderm (Cambray and Wilson, 2002) when placed in the right environment. Serial transplantations from the tailbud in chick into the primitive streak of earlier embryos also demonstrated their ability to repopulate the embryonic axis in this species, and their ability to re-set their hox expression based on their new environment (McGrew et al., 2008). Taken together, these studies reveal that the growth of spinal cord of amniote embryos is driven by a bipotent progenitor cell population located in the tail of the embryo and in a stage dependent manner. An argument can be built to link this population for the T/Bra, Sox2 expressing cells in the epiblast.

## Locating the neuromesodermal progenitor pool reveals multiple NMp populations

While the retrospective lineage experiments identify NMps as a stem cell population in the posterior region of the developing embryo, knowledge of exactly where this population resides requires precise fate mapping. In addition, it is important to ascertain whether a single NMp population gives rises to all of the axial structures or whether, as hinted at above, there exists multiple NMp populations that generate different medio-lateral compartments of the spinal cord and mesoderm.

The node is a prominent structure that appears just anterior to the primitive streak towards the end of gastrulation (Fig. 2). Fate mapping of this dynamic structure and its surroundings in the chick has shown a rostral-to-caudal landscaping in cell fate potential that ultimately end up in a medial-to-lateral position in the mesoderm once they have undergone an epithelial-to-mesenchymal transition and migrated to form the mesoderm (Freitas et al., 2001; Iimura et al., 2007; Psychoyos and Stern, 1996). First hints of the existence and location of NMps came from the injection of lysinated rhodamine dextran to cells in the node region of chicken embryos that generated extended clones with a dual contribution to both the floor plate and spinal cord (Selleck and Stern, 1991). This study also revealed that the node area is regionalized with respect to its contribution to mesodermal tissue: more rostral and medial positions give rise to clones of cells that are restricted to the notochord, more lateral labels in the node generate cells fated towards the medial aspect of the paraxial mesoderm and labels located both caudally and laterally to the node generate cells fated towards the lateral aspect of the somites (Selleck and Stern, 1991). Cells fated towards both floorplate and notochord are located within Hensen’s node (Selleck and Stern, 1991), whereas cells fated to both paraxial mesoderm and more lateral aspects of the spinal cord are located in regions both lateral and caudal to the node (Brown and Storey, 2000). This distinction between floorplate spinal cord progenitors in the node and lateral wall spinal cord progenitors being located in regions being located caudo-laterally to the node has also been observed with the use of quail-chick chimeras (Catala et al., 1996).

**Figure 2:**
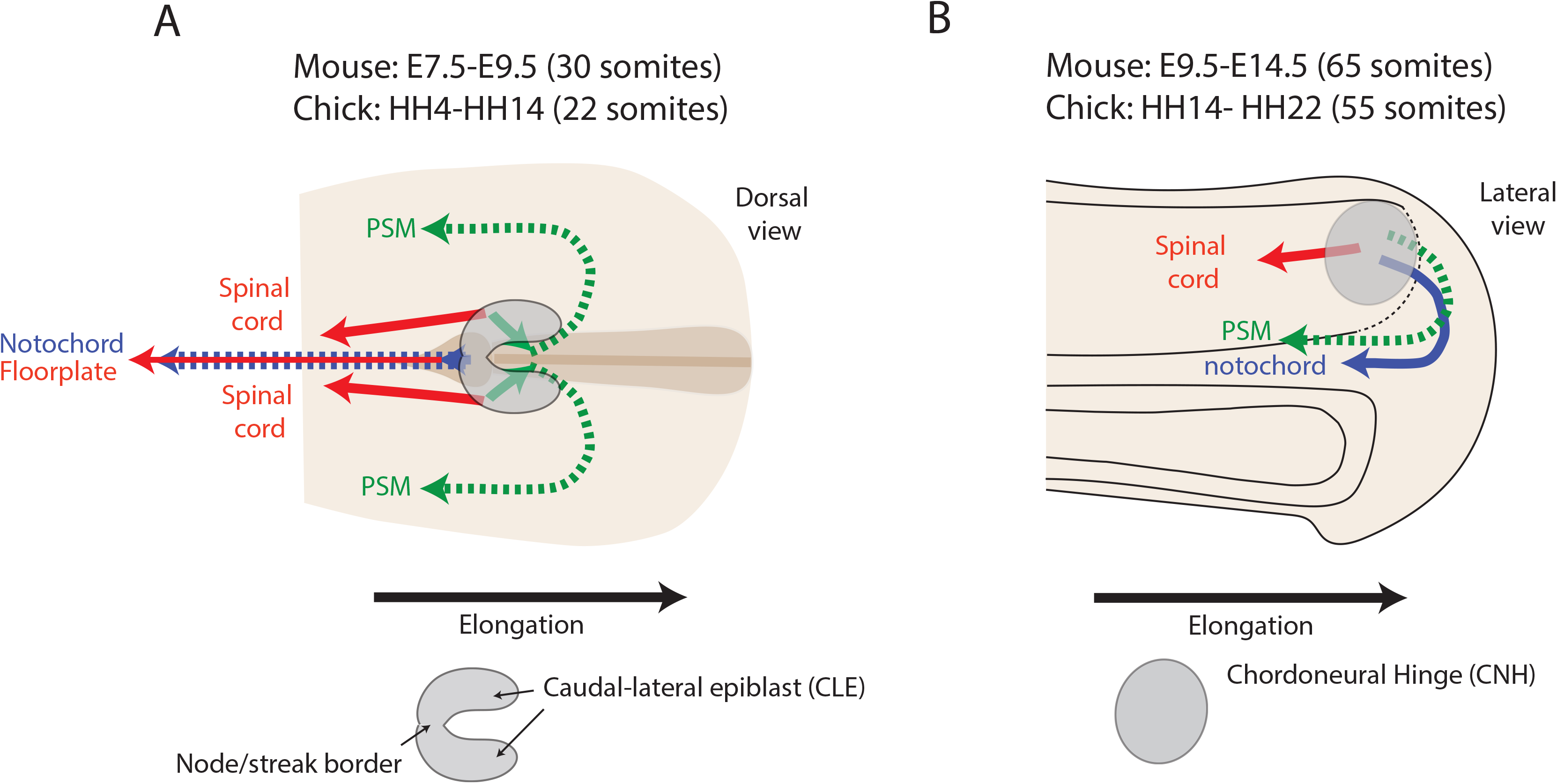
Locations of neuromesodermal progenitors at early and late stages of amniote development. Neuromesodermal progenitor territories are outline in grey, dotted lines represent cells that move out of the plane of the diagrammatic section. A) At early stages (after the completion of primary gastrulation and before the formation of the tailbud), the region surrounding and including the node generate cells fated to become both spinal cord and mesodermal cells. Within the node, cells generate the notochord and floorplate. Posterior and lateral to this, cells generate the medial aspect of the spinal cord and somites. B) Upon formation of the tailbud, a region called the chordoneural hinge contains progenitors of the spinal cord (red), Pre-somitic mesoderm (PSM; green) and notochord (blue).

Detailed transplantation experiments in mice reveal that while the region of epiblast caudal and lateral to the node (CLE) and the node-streak border (NSB) can give rise to both neural and mesodermal derivatives, regions more caudal to this cells are solely fated towards mesoderm (Cambray and Wilson, 2007, 2002; Wymeersch et al., 2016). Taken together, these studies suggest cell fates are highly regionalised around the node in amniotes (Figure 2). Importantly however, heterotopic transplantations from more caudal regions into the NMp region are able to generate dual neural/mesoderm fated cells (Cambray and Wilson, 2007; Wymeersch et al., 2016). This is limited to the region just caudal and lateral to the node, as transplants of more caudal regions are restricted to mesoderm fate (Wymeersch et al., 2016). Furthermore, grafts of primitive streak tissue from older mouse embryos into earlier staged host continue to contribute to more anterior somites (Tam and Tan, 1992). Therefore, whilst prospective mesoderm tissue maintain a certain degree of plasticity in their fates until late stages of development, the ability to generate both neural and mesodermal derivatives is regionalised. Taken together, these data suggest that there are at least two populations of NMps cells, defined as cells that will give rise to both mesoderm and caudal tissue, in amniote embryos. One gives rise to the notochord (axial mesoderm) and floorplate (ventral SC), while a second population generates the medio-lateral aspect of paraxial mesoderm and the spinal cord (Figure 2A). In amniotes, these cell populations undergo temporally restricted self-renewal to generate a large proportion of the posterior body axis, and is in line with the relative small region that gives rise to the spinal cord in gastrula stage fate maps (Figure 1).

## Clonal continuity of NMp populations along the anterior-posterior axis

The fate mapping experiments described above suggest that the NMps generate central nervous tissue from approximately the base of the hindbrain, to what posterior extent do they contribute to the spinal cord? Upon formation of the tailbud, prospective paraxial mesodermal tissue folds underneath the posterior end of the forming secondary neural tube, with the very posterior end of this attached to the posterior aspect of the notochord (Knezevic et al., 1998). This conjoined structure is termed the chordoneural hinge (CNH; Figure 2B). Fate mapping with the use of chick with quail-chick chimeras showed that the chordoneural hinge (CNH) moves in a rostral-caudal direction and gives rise to the floor plate of the lumbo-sacral-caudal neural tube. Furthermore, a region within the tailbud that is medial and caudal gives rise to cells that diverge laterally and ultimately contribute to both the somites and to the spinal cord, it is likely that this is the region in which NMps arise in the chick (Catala et al., 1996). Therefore, while these CNH NMps apparently contribute to the posterior-most extent of the spinal cord, a key outstanding question is whether the NMp population that is apparent in the CLE and NSB at early stages has clonal continuity with cells in the CNH upon tailbud formation.

The ability of CNH derived grafts to generate both neural and mesodermal tissue of the whole axis when transplanted into early stage hosts strongly suggests that they maintain at least some of the properties of early stage NMp populations surrounding the node (Cambray and Wilson, 2007). In addition, homotopic grafts of E8.5 CLE generate descendants within the CNH (Cambray and Wilson, 2002) and gene expression studies reveal the existence of cells coexpressing T/Bra and Sox2 in this region and suggest a degree of topographical continuity between the CLE and the CNH (Cambray and Wilson, 2002; Wymeersch et al., 2016). Importantly, a similar continuity between early and late NMp populations has also been observed through heterochronic grafts in the chick (McGrew et al., 2008). Furthermore, the retrospective clonal analysis study of Tzouanacou and colleagues (Tzouanacou et al., 2009) showed that there are bipotent clones that span the trunk/tail transition, thus supporting clonal continuity of the CLE/NSB and CNH progenitor pools. However, while the increase in the frequency of bipotent clones is indicative of an expansion of this population up until tailbud formation, many clones arrest at the level of the posterior trunk, suggesting that from tailbud stages onwards there is an overall depletion of the NMp pool from this stage onwards (and see Wymeersch et al., 2016). Furthermore, live imaging of notochord development in the mouse reveal distinct cell behaviours that lead to the elongation of the notochord depending on axial level (Yamanaka et al., 2007). Therefore, while clonal analysis and transplantation studies do suggest a degree of clonal continuity NMp populations throughout the process of axial elongation in amniotes, further live imaging studies are required to understand how dynamic cell behaviours of NMps are altered at each stage of this complex process. Taken together, these lineae tracing studies suggest that multiple populations of NMps exist, that alter in both their medio-lateral and anterior-posterior contribution to the elongating body axis.

Understanding the relative contributions of these progenitor pools, and how this differs between vertebrate models is important in order to gain a complete picture of spinal cord development in vertebrates.

## Neuromesodermal progenitors in anamniotes: plasticity in the absence of self-renewal

It has been suggested that zebrafish embryos harbour a bipotent self-renewing population of NMps (Martin and Kimelman, 2012). Indeed, challenging single cells with a range of cell-autonomous signal perturbations has clearly shown that distinct populations of cells retain the ability to generate multiple germ layer derivatives into somitogenesis stages (Martin and Kimelman, 2012, 2008; Row et al., 2016). However, fate maps in zebrafish and Xenopus embryos show that, at the gastrula-stage, there already exists a significant region of tissue that is fated to become spinal cord (Figure 1). Therefore, it is not clear the degree to which a self-renewing pool of progenitor cells is required to generate the posterior body axis of these organisms.

Single cell labelling within the shield region (functionally equivalent of the amniote node) revealed a mixed population of floorplate and notochord progenitors with no dual neuronal and mesodermal fated cells and little evidence for long term self-renewing progenitors (Shih and Fraser, 1995). Furthermore, the clones produced from each labelled cell are not long clones like those seen in amniotes (Brown and Storey, 2000; Selleck and Stern, 1991; Tzouanacou et al., 2009) but rather short clones in the range of 1-8 cells spreading over 1-6 somites in length (Shih and Fraser, 1995). In addition, the single cell grafts into the marginal zone by Martin and Kimelman (2012) showed only very few cells giving rise to dual neural and mesodermal derivatives in the wild-type situation. In order to generate a progenitor/stem cell mode of growth, a population must divide rapidly enough to generate self-renewal during the process of axis elongation. At gastrula stages, clones of photolabelled cells that enter directly into the trunk spinal cord do so only with very low proliferation rate of a mean of less than 0.1 cells per hour (Steventon et al., 2016). Upon formation of the tailbud, this drops even further for tailbud progenitors (Bouldin et al., 2014; Kanki and Ho, 1997; Steventon et al., 2016). Furthermore, blocking of cell division at late gastrula stages has only a minor impact on axis elongation as a whole (Riley et al., 2010; Zhang et al., 2008) and single cell labels in the tailbud studied do not support the existence of bipotential neural/mesodermal progenitors (Kanki and Ho, 1997).

The embryonic shield in zebrafish appears to be transient region as reflected by the short length clones produced from single cell injections (Shih and Fraser, 1995) and similar restricted clonal labelling is observed in the Xenopus gastrula (Keller, 1975). This continuous transition of cells through the anamniote organiser region is also seen during primitive streak stages in the chick up until stage 4 (Joubin and Stern, 1999). Therefore, during primary gastrulation in both anamniotes and amniotes the node/organiser/shield region represents a transient structure through which cells pass and then generate axial structures by convergence and extension. However, in mice and chick this is followed by a second phase, associated with the emergence of the node, involving the expansion of NMp populations to generate axial structures until the point at which the posterior neuropore closes and the tailbud is formed. It is this second stage that is absent in anamniotes, that instead go directly from primary gastrulation to the formation of the tailbud.

The tailbud of frog embryos have a distinct CNH and DiI labelling of the blastoporal lip at stage 13 in Xenopus leavis results in the labelling of CNH at tailbud stages (Gont et al., 1993). This suggests that, like the node labels in mouse and chick, the organiser later gives rise to derivatives that form the CDH, and later the floor plate and notochord. At the completion of gastrulation blastopore labels generate the tailbud and its derivatives but not more anterior structures (Gont et al., 1993). Fate mapping of the tail-forming region at late neural stages in Xenopus revealed that much of the tail region is not generated from the tailbud but rather from a posterior displacement of trunk tissue into the tail region (Tucker and Slack, 1995). This is also the case in zebrafish embryos, where a continued growth of the spinal cord in already segmented region of the body axis results in a posterior displacement of this tissue relative to the pre-somitic mesoderm that is undergoing addition of cells by convergence and extension movements in the tailbud (Steventon et al., 2016). Labelling of small groups of 3-4 cells within the tailbud of late stage Xenopus embryos result in the labelling of both neural tube, notochord and somitic tissue. While these results do not demonstrate the production of multiple germ layer derivatives from single cells, they do suggest that neural and mesodermal progenitors lie in close proximity until late stages within the CNH (Davis and Kirschner, 2000).

These differences in the modes of growth of NMp cells between amniotes and anamniotes may be underlined by the vast differences in the degree of volumetric growth that occurs concomitantly with posterior body elongation. Mice embryos undergo an approximate 65-fold increase in their posterior body volume from the tailbud stage to the completion of somitogenesis, which is much higher than the 4-fold increase that is observed in zebrafish embryos (Steventon et al., 2016). This difference is most likely due to differences in the degree of overall growth that occurs concomitantly with embryogenesis in internally vs. externally developing embryos. Despite these differences in the cellular processes generating the spinal cord, the experiments of Kimelman and colleagues clearly demonstrate a plasticity in germ layer restrictions within tailbud progenitors of zebrafish embryos (Martin and Kimelman, 2012, 2008; Row et al., 2016). Therefore, while NMps do not undergo overt self-renewal in this embryo, they do seem to exhibit a bipotent neural/mesodermal cell state.

## Cell populations and continuity during vertebrate axial elongation

Taken together, the studies to date demonstrate that central nervous system progenitors exhibit a different modes of elongation, depending on their ultimate position along the anterior-posterior axis. Firstly, convergence and extension based movements that generate tissue elongation by cell rearrangements. Secondly, stem cell growth from an expanding pool of NMps situated in the node region, and finally growth from a depleting NMp pools located within the CNH. The relative contribution of these different growth modes to the axis differs widely between the four key vertebrate models that have been studies (Figure 3). While the dispersive mode of growth that is characteristic of the convergence and extension movements of gastrulation generate the forebrain, midbrain and hindbrain of both mouse and chick embryos, these cellular behaviours extend much further to cover the trunk region of both zebrafish and Xenopus embryos (Figure 3; purple lines). Upon completion of primary gastrulation in amniotes, cells within and adjacent to the node proliferate and thereby provide a self-renewing pool of bipotent progenitors that generate the spinal cord up until the closure of the posterior notochord (Figure 3; blue lines). In the absence of overt self-renewal, this process does not occur in anamniotes, as primary gastrulation is followed directly by the closure of the blastopore and the formation of the tailbud. Upon tailbud formation in all species examined, the neuromesodermal progenitor pool becomes located within the CDH from which the tail spinal cord is generated (Fig. 3; green lines). From this stage onwards, the NMp population becomes continually depleted until the end of somitogenesis. Thus from the cellular point of view, the main difference between anamniote and amniotes is the dynamics of the NMps that will give rise to the SC; progressive depletion of a large pool generated before gastrulation (anamniotes) or generation of the pool through the amplification of a small progenitor pool (amniotes). The emergence of the CNH signals a process that is common to both in which the NMp pool is slowly used up through processes of convergence-extension. The vast differences in modes of growth of spinal cord progenitors offers a challenging and fascinating problem to developmental biologists: how do seemingly conserved signal and gene regulatory networks map on to these vastly differing geometries and cellular behaviours? In the next section, we will attempt to touch on this problem by reviewing what is known about the molecular mechanisms that are important for balancing the maintenance and differentiation of NMps.

**Figure 3:**
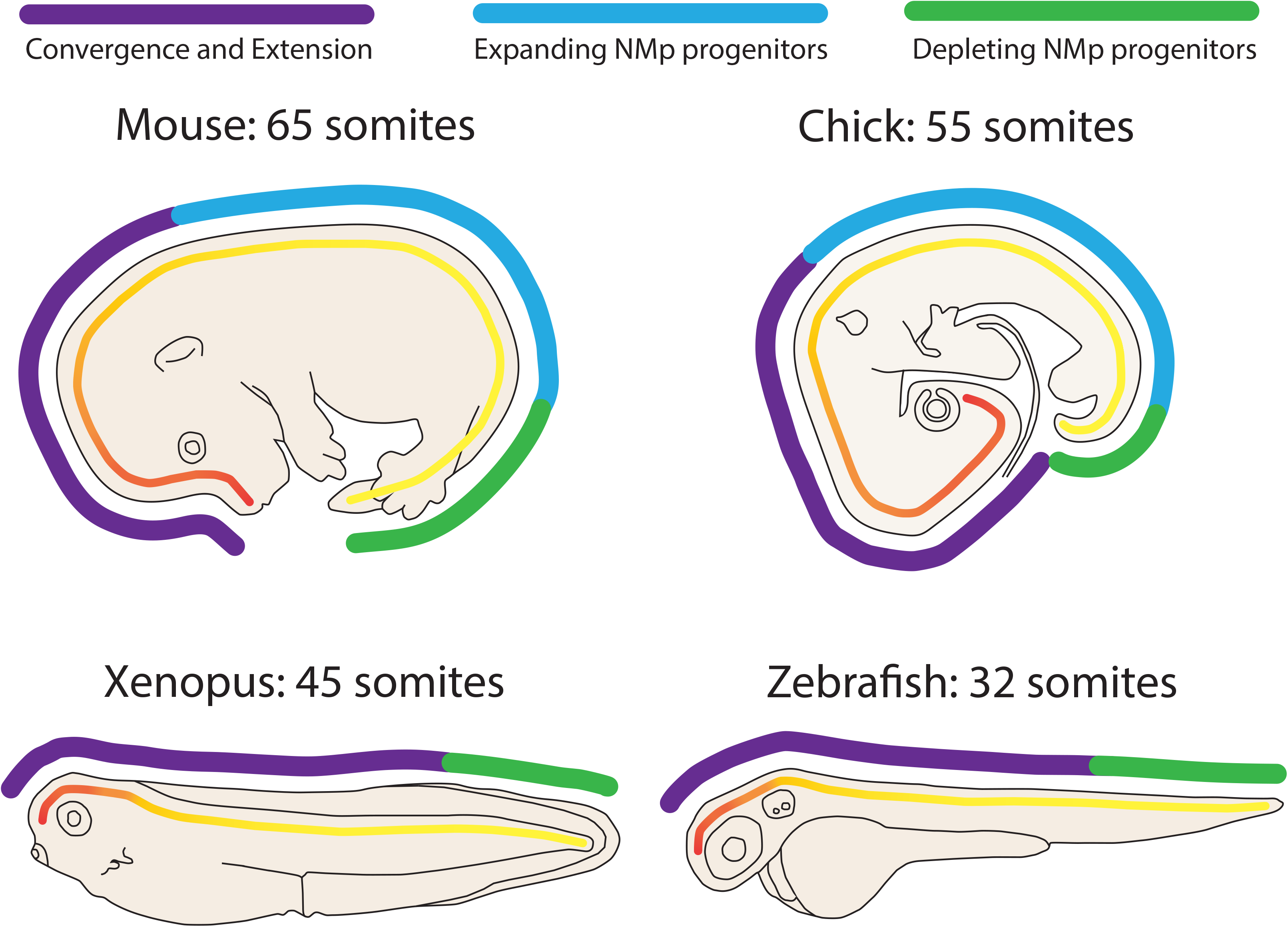
Differential contribution of cellular behaviours to the generation of the spinal cord in vertebrates. Three distinct cellular behaviours can be attributed to the generation of the spinal cord in vertebrates. To summarise the differences outlined between the contributions of these behaviours between the amniote models mouse and chick (upper row) to those in the anamniote *Xenopus* and zebrafish (lower row), these have been mapped onto the late stage embryos as shown. Convergence and extension (purple bar) generated the brain region in all four models. However, the expanding neuromesodermal populations (NMps) situated by the node (blue bar) in amniotes is absent ananmiotes that transition directly from gastrulation to the formation of the tailbud population of depleting NMps (green bar).

## Balancing self-renewal and differentiation of NMps

An accepted molecular definition of NMps is as cells that coexpress Sox2 and T/Bra and thus express the possibility of becoming neural and mesodermal. Much of the control of axial elongation relies on the maintenance of cells co-expressing these transcription factors and is tightly linked to the onset and activity of members of the Cdx and Hox families of transcriptional regulators (Young and Deschamps, 2009). Cdx genes encode homeobox containing transcription factors which are expressed in the primitive streak during gastrulation and then become localized to the tailbud of the growing embryo in a manner similar to T/Bra. In the mouse there are three Cdx paralogues (1,2 and 4) which exhibit functional redundancy (Chawengsaksophak et al., 2004; Subramanian et al., 1995; van de Ven et al., 2011; van den Akker et al., 2002). Postimplantation removal of the three paralogues reveals a clear deletion of the axial component of the body plan posterior to the occipital region (somite 5), that is associated with a loss of the NMp population (van Rooijen et al., 2012). Importantly however, a main role of Cdx genes appears to be to establish an appropriate signalling environment for axial extension, as cells deficient in Cdx can correctly participate in axial elongation when placed in a wild-type environment (Bialecka et al., 2010). Indeed, genetic analysis suggests that Cdx proteins implement their activity in a complex manner by regulating the expression of posterior Hox genes (Hox5-Hox13) and non Hox targets such as ligands for FGF (4 and 5) and Wnt (8a and 3a) signalling. This is demonstrated by the observation that Hox genes from the trunk region (Hoxa5 and Hoxb8), expressed under the control of Cdx2, will rescue the Cdx mutant phenotype as does expression of an activated form of the Wnt/ß-catenin effector LEF1 (Young et al., 2009) or exposure of the Cdx mutant embryos to FGF signalling (van Rooijen et al., 2012). A connection between Hox expression and axial extension is further emphasized by the observation that expression of Hoxb13 under the control of Cdx2 truncates the extension of the body mimicking the Cdx mutant phenotype (Young et al., 2009). In addition, mice homozygous for Hoxb13 show an overgrowth of tail structures (Economides et al., 2003).

The connection between Cdx and Hox gene function and axial elongation is intriguing. Hox proteins are not specific regulators of defined processes but rather transcription factors that provide context to other regulators (Noordermeer and Duboule, 2013). What those events are in the case of axial extension is not clear but there is an intriguing correlation between the onset of expression of Hox5-11 and the proliferation of NMps (Forlani et al., 2003; Wymeersch et al., 2016). This suggests that trunk Hox genes could contribute to the process of axial elongation by creating responsiveness of the tail bud area to proliferative signals between E7.5 and E10 and to the termination of this process by Hox13 paralogs. In addition to controlling the population dynamics of NMps, the Hox code has also been shown to be important in controlling the timing of cell ingression into the primitive streak by interacting with Wnt signalling, and ultimately the specification of mesoderm (Denans et al., 2015; Iimura and Pourquié, 2006). Some of these effects might be non-autonomous or niche related and might underlie the observation that heterochronic transplants of the CNH allow for long term proliferation of the NMp region (Cambray and Wilson, 2007, 2002; McGrew et al., 2008). Indeed, cells mutant for Cdx can be rescued when transplanted into a wild-type environment (Bialecka et al., 2010). A role for colinear Hox gene activation as a timer for commitment to anterior-posterior identities has been proposed in the frog embryos as cells from non-organiser regions are brought into close proximity with Spemann’s organiser by convergence and extension movements (Wacker et al., 2004).

In addition to the role of Hox proteins in providing a context for the proliferation and differentiation of the NMps, there are genetic arguments to support a role for interactions between FGF, Wnt and RA signalling modulating both the balance of self-renewal and differentiation of NMps (Mathis et al., 2001; Nordstrom et al., 2002; 2006; (Abu-Abed et al., 2001; Akai et al., 2005; Cunningham et al., 2015; Diez del Corral et al., 2002; Kumar and Duester, 2014; Olivera-Martinez and Storey, 2007; Olivera-Martinez et al., 2012a, 2012b; Ribes et al., 2009). However, understanding the relationship between Cdx, Hox, FGF, Wnt and RA signalling in the activity of the NMps is challenging and requires an understanding of their temporal relationships. For example Cdx and Hox expression is initiated independently during gastrulation, with an important input of Wnt signalling (Forlani et al., 2003) but by the time the NMp population is established at E8.0, hierarchical relationships emerges: Hox gene expression falls under the control of Cdx and a number of reciprocal feedback loops are established between Cdx, FGF4, 8 and Wnt3a and 8a that promote the proliferation of the NMp initial pool and coordinate their differentiation (Fig. 4). While it is possible that Cdx proteins have a cell autonomous role in the establishment of the NMps, the rescue of Cdx mutants with FGF and Wnt signalling suggests that a most important role of Cdx is to create a niche for the NMps and probably coordinate its activity with the anterior posterior level by regulation of the Hox genes.

**Figure 4:**
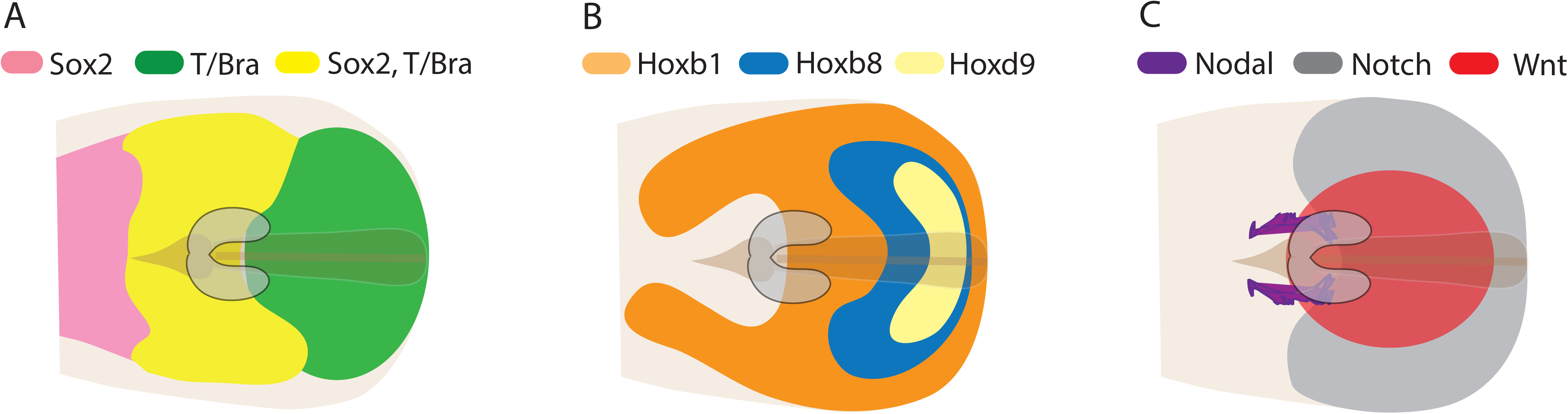
Gene expression domains around the node during the NMp expansion phase. Simplified diagrams of the node/streak region of amniote embryos are shown with anterior to the right and posterior to the left. Grey horse-shoe domain demarcates approximate position of the NMps. A) Expression of sox2 and bracyury/T, B) Expression of Hoxb1,8,9 and C) Appoximate activity maps for Nodal, Notch and Wnt.

The function of the different signaling pathways in the development of the spinal cord has been studied by the use of gain and loss of function constructs into chick embryos. By electroporating cells with a dominant negative FGF receptor construct together with a lineage tracer, a clear role for FGF in the maintenance of NMp self-renewal has been demonstrated as cells defective in FGF reception undergo a precocious differentiation and exit from the progenitor pool (Mathis et al., 2001). Furthermore, localised exposure to increased FGF levels results in the inhibition of neuronal differentiation and the maintenance of progenitors (Diez del Corral et al., 2002) whose proliferation is driven by Notch signalling downstream of FGF (Akai et al., 2005). FGF signalling is balanced by a somite-derived retinoic acid (RA) signal that promotes the differentiation of neuronal differentiation by forming an opposing anterior-posterior gradient of signalling (del Corral and Storey, 2004; Diez del Corral et al., 2003). Whilst important for maintaining progenitors through interactions with FGF signalling, Wnt signalling also facilities the switch to differentiation by promoting retionic acid receptor expression once FGF signalling has been attenuated (Olivera-Martinez and Storey, 2007). Similar interactions have been observed in the mouse, where RA has been shown to provide an element to restrict the tail bud region, where Cyp26A1, under the control of Wnt and FGF, keeps it free from RA (Young et al., 2009). These interactions provide a mechanism by which posterior elongation comes to a timely end, as a progressive loss of FGF signalling leads to an increase in retinoid signalling, thereby switching of genes responsible for the maintenance of NMps within the chordo-neural hinge and an increase in cell death (Olivera-Martinez et al., 2012b). It is likely that these events are coordinated by the expression of the posterior Hox genes.

Taken together, these observations can be incorporated into a tissue level model for axial elongation in amniote embryos in which NMps arise at the end of gastrulation at the same time as and around the node (Figure 5). Subsequently, at around E8.0-E9.0 in the mouse, NMps undergo an expansion that is dependent upon canonical Wnt signalling (Wymeersch et al., 2016). Thereafter, FGF, Wnt, Notch and RA signalling interact to generate a complex signalling networks that determines the precise balance of proliferation, differentiation and apoptosis to generate a correctly proportioned body axis (see Fig 5 for details). Recent experiments have shown that Wnt signalling plays a main role in driving the amplification of the NMps. A significant component of the NMps is identified by a specific enhancer of the Sox2 gene, the N1 enhancer, which becomes active at the time of emergence of the NMps driven by a combination of Wnt and FGF signalling (Takemoto et al., 2006). Interestingly, this enhancer is absent in the zebrafish sox2 gene, although the mechanistic consequence of this is (Kamachi et al., 2009). Interestingly, Wnt, FGF and retinoic acid have all been suggested to be candidates for posteriorizing factors that are important for spinal cord formation during gastrulation in *Xenopus laevis* (Bang et al., 1999; Holowacz and Sokol, 1999; Isaacs, 1997; McGrew et al., 1995; Niehrs, 2001; Simeone et al., 1990; Villanueva et al., 2002). How these upstream regulatory processes have been co-opted and adapted to the differing cellular behaviors that drive axial elongation in amniotes is an open question.

**Figure 5:**
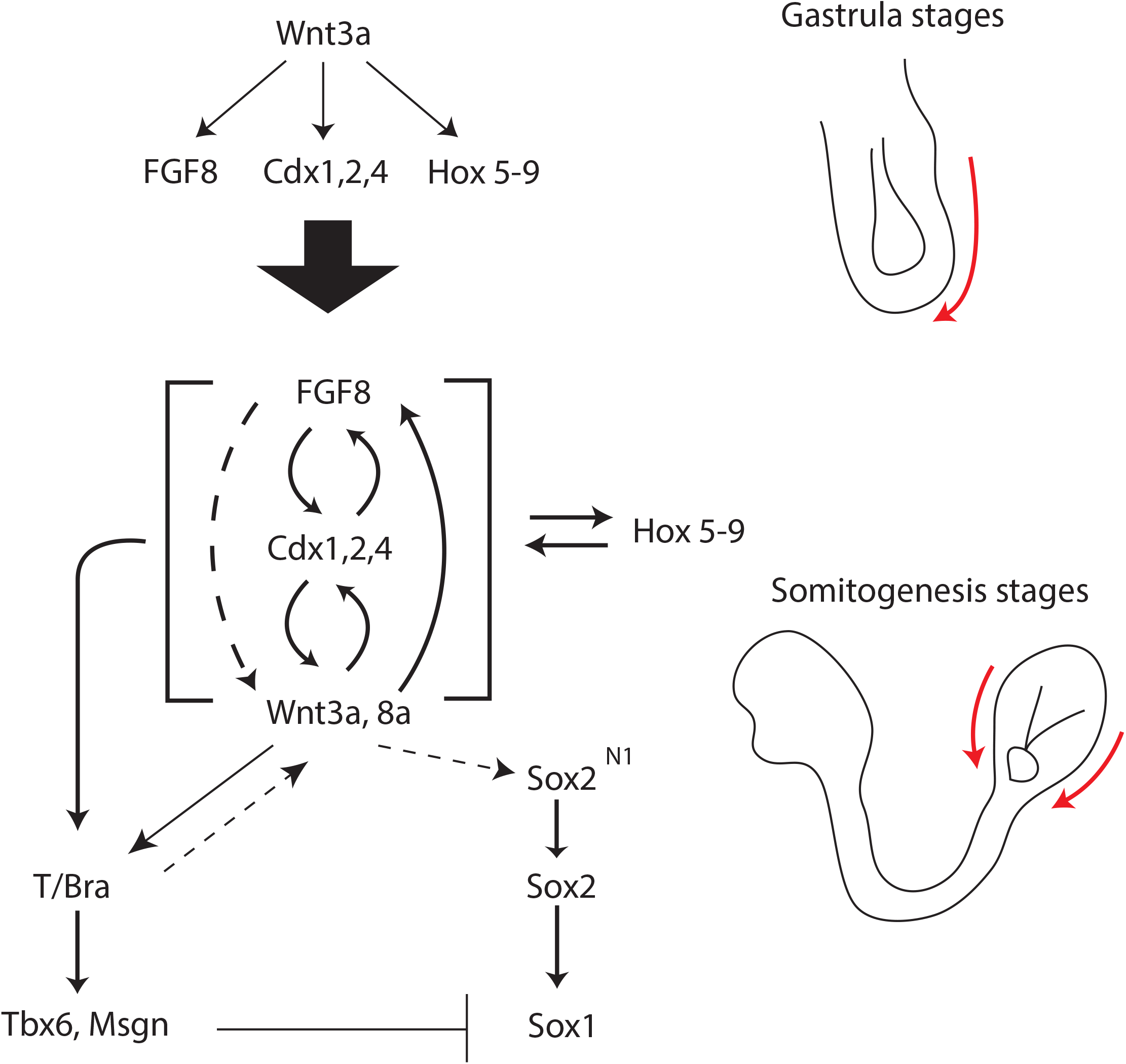
Gene regulatory networks during the establishment and expansion of NMps. At gastrula stages (upper panel), the NMp niche is established downstream of Wnt signaling by the activation of FGF, Cdx2 and Hox 5-9. Subsequently, an autoregulatory loop is established between these factors that maintain the NMp population through somitogenesis stages (in brackets). Finally, competing interactions between early neural and mesodermal markers determine cell fate choice as cells exit the NMp pool (lower panel).

## Balancing neural vs. mesodermal cell fates

In addition to balancing the cell population dynamics of NMp proliferation, extracellular signalling networks must also interact with master regulators of both neural and mesodermal gene regulatory networks to precisely balance cell fate. Unlike the NMp expansion, this is likely to be conserved in amniotes and anamniotes. Experiments in zebrafish have been particularly informative in beginning to tease out the mechanisms underlying this decision with the use of single cell transplants and temporally-controlled functional experiments (Martin and Kimelman, 2012; Row et al., 2016). Upon transplantation into the marginal zone at the embryonic shield stage, cells are normally fated to enter into the paraxial mesoderm, but inhibition of Wnt signalling with a dominant-negative TCF construct results in redirecting the fate of these cells to a neuronal lineage (Martin and Kimelman, 2012). Similarly, inhibition of Wnt signaling in a population of midline cells that either give rise to the notochord, floorplate or hypochord, biases cells to the neural lineage over both notochord and hypochord. Activation of Wnt signalling has the reverse affect (Row et al., 2016)

While these data clearly demonstrate that, in the zebrafish, cells retain a degree of plasticity in their ability to be pushed towards towards neural or mesodermal cell fates, they do not show that axial elongation is driven by a self-renewing pool of stem cells as it occurs in the mice and chick (see above). However, the differential requirement for the differentiation process appears to be conserved and in mice loss of Wnt signaling leads to a secondary neural tube and a depletion of the mesoderm (Nowotschin et al., 2012) whereas increases in Wnt signalling augment the mesodermal pool but do not affect neural development (Garriock et al., 2015). There may be some specific differences in the role of Wnt signaling in the specification of mesoderm however, as in zebrafish it appears to act to promote mesodermal specification the expense of neural (Martin and Kimelman, 2012; Row et al., 2016).

The targets of these signalling events are likely to be the expression of Sox2 and T/Bra. The levels of these transcription factors matter and it has been shown recently how the NMp population resides in a region with low levels of T/Bra expression within uniformly low levels of Sox2 (Wymeersch et al., 2016). The co-expression of Sox2 and T/Bra as a substrate for the decision also appears to be the case in the zebrafish tailbud (Martin and Kimelman, 2012).

In the context of our discussion, we would define an NMp in both amniotes and anamniotes, as an epiblast derived cell that undertakes a fate decision process between neural and mesodermal fates. The difference between the two groups is that in amniotes NMps undergo an amplification step which is absent in amniotes, likely a consequence of posterior body elongation occurring concomitantly with growth (Steventon et al., 2016). Importantly, the underlying structure of signal and gene regulatory networks that act to balance NMp population dynamics and their decision to generate either neural or mesodermal cell fates are largely conserved (Martin and Kimelman, 2009). To understand how these conserved regulatory networks map to such vastly different cellular substrata, we must take a dynamical systems approach. In other words, we have to understand how slight changes in regulatory inputs can act to alter the rates of cell fate transitions and population dynamics between experimental systems.

### The evo-engineering of axial elongation: a dynamical systems view of the NMps

As it is often the case with cell states, NMps are defined by the coexpression of some transcription factors, in this case T/Bra and Sox2 with, sometimes Nkx1.2. However, we would like to suggest that rather than a specific state, NMps represent what we have termed a ‘transition state’ (Arias and Hayward, 2006; Muñoz-Descalzo et al., 2012): a metastable arrangement of gene regulatory networks during a binary cell fate decision. This arrangement allows a cell to express simultaneously genes associated with the fates involved in the decision thus priming both fates and allowing a form of bet hedging based on fluctuations in the levels of the alternative fates. At the level of a population, the transition state is characterized by heterogeneities in gene expression. In the context of NMps, the existing observations allow us to suggest the following sequence of events.

Cells enter the transition state from the epiblast, where they already express *Sox2*. Under the control of Wnt/ß-catenin signallling at the NSB, they reduce the levels of *Sox2* and activate expression of *T/Bra* thus creating the well characterized NMp signature in which cells co-express *T/Bra* and *Sox2* and many genes under the control of these two transcription factors which thus creates a substrate for differentiation at the level of individual cells. It is not known the relevance of the levels of Sox2 but there is evidence that the levels of T/Bra are low in NMps and high in mesodermal progenitors (Turner et al., 2014). We would surmise that the same applies to the levels of Sox2: low in NMps and high in neural progenitors. Thus, increases in T/Bra expression will lead to activation of **Tbx6**, **Mesogenin** and other genes associated with mesoderm development, while increases in Sox2 expression will lead to Sox1 and neural fate associated genes (Olivera-Martinez et al., 2014).

In the transition state cells maintain low varying levels of the different fate associated genes which allows the decision and this state would be represented by cells with low, and perhaps fluctuating, levels of both Sox2 and T/Bra. If during the time that it takes to make the decision cells in this state can divide, this will lead to an amplification of the transition state and it is in this sense that they will create a long-term progenitor. The levels of Sox2 and T/Bra are likely to be controlled by the levels of Wnt and FGF signalling (Figure 6) and fluctuations and gradients of these signals will create spatial landscapes that will determine the probability that a given cell remains in the transition state or differentiates on the basis of its position. The deployment of elements of Notch signaling in the tail bud suggests that they are also involved in this decision (Figure 6) and, indeed, there is evidence that Wnt and Notch signaling play interacting and opposing roles in the maintenance and resolution of the transition state (Arias and Hayward, 2006; Muñoz-Descalzo et al., 2012) and this might also be the case here (Figure 6).

**Figure 6:**
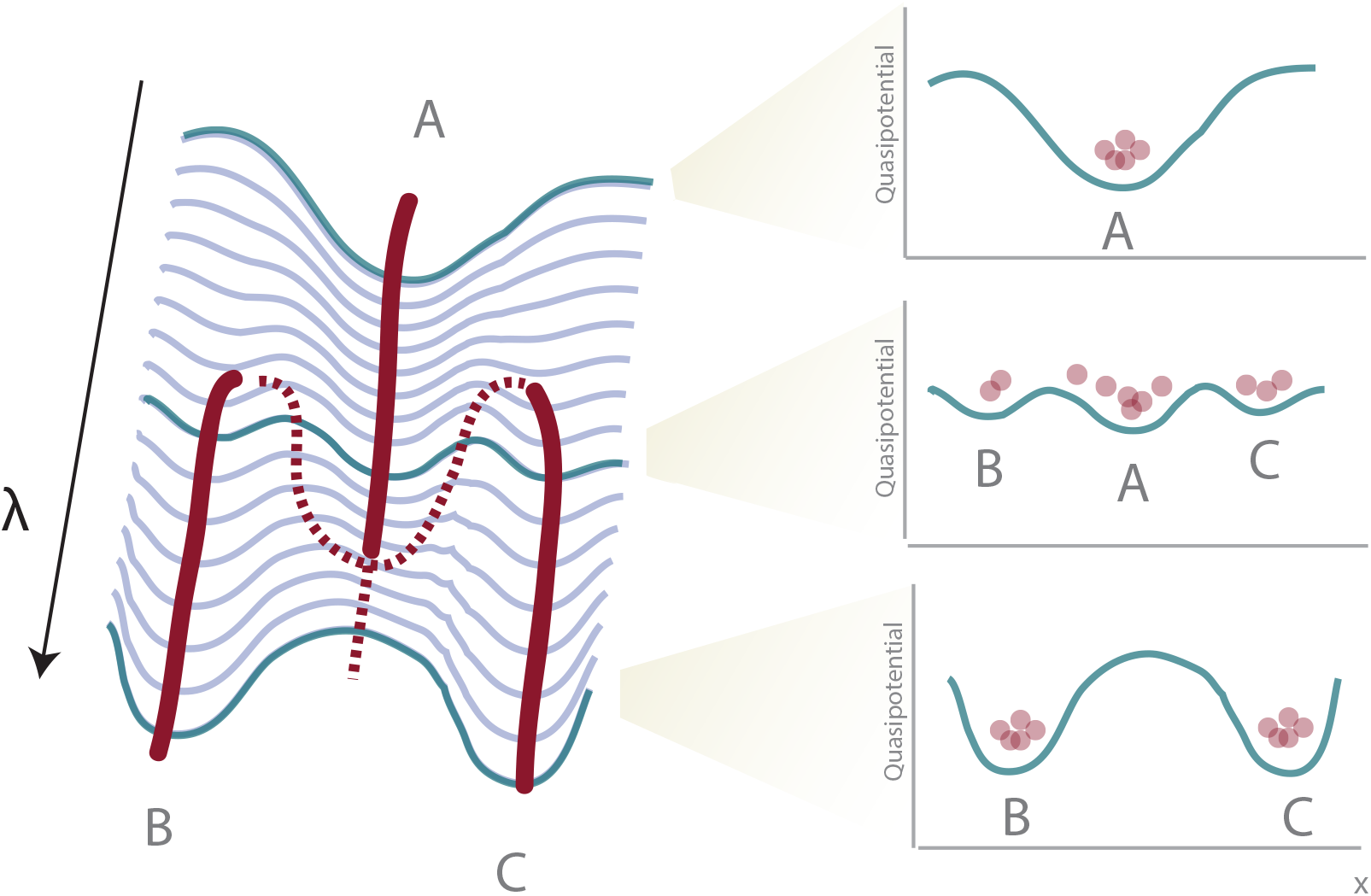
NMps as a transition state can be understood by a subcritial pitchfork bifurcation model. Understanding neuromesdermal progenitors in terms of a transition state (A) from which neural (B) or mesodermal (C) stable states emerge can be modelled in terms of a subcritical pitchfork bifurcation model. On the left shows that this can be represented in the form of a Waddington landscape with cells either proceeding in the transition state (A) and thereby in a neuromesodermal state or resolving into either of the two stable cell fates (B,C).

We propose that the transition state is common to both amniotes and anamniotes and only the ‘self renewal’ changes between the two i.e. if cells can divide while remaining in the transition state, they will be amplified. It is likely that this is the main difference between amniotes and anamniotes. Inspection of the relationship between the landscape and the NMp population reveals some of the molecular elements governing the TS (see Figure 5).

As cells exit the transition state, they undergo fate decisions that can be modelled in terms of dynamical systems theory as resulting from the dynamics of interactions between elements of GRNs. A formal representation of these interactions leads to a set of equations whose solutions depend significantly on the value of the parameters of the networks e.g. the rate constant of the interactions between transcription factors and target genes. Analysis of how the solution of the equations depends on the parameters of the system reveals the existence of situations in which the system can choose between more than one stable solution. In the context of developmental systems, this would be a point of fate choice: the point at which the system evolves different solutions according to its parameter is called a ‘bifurcation’, because the solutions ‘bifurcate’(Ferrell, 2012)-refs-).

There are different kinds of bifurcation and some of them can be adapted to developmental processes. Thus, cell fate decisions in the famed Waddington landscape have often been related to a *supercritical pitchfork bifurcation* in which a stable state (fate) gives rise to two alternative ones. The strengths and weaknesses of this straightforward view of Waddington landscape have been discussed and do not fit many of the features of cell fate decisions in development (Ferrell, 2012). A specific problem with the pitchfork bifurcation is that, while being an intuitive representation of a biological process, it cannot account adequately for the directionality of the fate decisions nor for the structure of the transition state. A different class of bifurcations, *subcritical pitchfork*, can be used to represent many of the features of a transition state. In this type of bifurcation, for a certain value of the parameters of the system, there is a solution in which a starting and two alternative states can coexist (Fig. 6; Huang et al., 2007). In the context of the NMps this would give rise to a state in which a cell would express in a dynamic manner markers of the epiblast as well as of the neural and mesodermal progenitors, as is observed. (Turner et al., 2014). In this representation, the driving force of the system resides in the Gene Regulatory Networks that promote the changes in state (the genes regulated by T/Bra and Sox2), and the parameters that control the stability of the states are the extracellular signals (Huang et al., 2007) in particular Wnt, FGF and Notch.

This formulation leads to a number of considerations which help understand the relationship between the NMps of amniote and anamniote embryos from an engineering point of view. The two most important features that distinguish the two groups are the self-renewal of the NMps and the stability of the NMp state. In a simplest model, increased self-renewal maybe due to an overall increase in proliferation across the body axis that is linked to body axis formation occurring concomitantly with growth (Steventon et al., 2016). In terms of differences in the stability of the NMp transition state, understanding this in terms of dynamical systems modelling may be of great help. The range of the instability is determined by the value of some critical parameter of the system that determines the stability of each of the states involved in the decision. Interactions between signalling and gene regulatory networks could be incorporated into a parameter ⌊, which would determine the extent of the transition state. For a certain critical value ⌊;c, there would be a transition state (amniotes) but as ⌊c –->0 a situation is reached, as in the anamniotes, in which the subcritical bifurcation becomes a pitchfork bifurcation. We appreciate that this is an oversimplification but we hope that this view will encourage a consideration of the decision on formal terms. This representation also allows for an understanding of the evolutionary changes of the nature of NMp population and highlights its control as understanding the molecular nature of the parameter ⌊c will provide insights into the mechanisms that underlie the evolutionary plasticity of the NMp state. We would surmise that the networks underlying the different states (epiblast, neural and mesodermal progenitors) involved in these decisions are conserved and that the difference lies in the control of the parameters that govern these two features.

## Conclusions and future directions

Despite the apparent conservation in molecular mechanisms that act to pattern the vertebrate neural axis at the cell population level, there are differences in the underlying cell behaviours that act to elongate the body axis at the same time as establishing the spinal cord. In order to understand how seemingly conserved signal and gene regulatory networks can act together with these different cell behaviours to generate body axes of varying proportions, we invoke the concept of a transition state in which NMps are trapped in the decision to either generate spinal cord or mesodermal derivatives. This is important, as it is the first step towards generating dynamical systems models of NMp self-renewal and differentiation that that have the potential to explain how the embryos creates a flexible patterning mechanism that can be mapped onto cellular substrates of differing geometries. We propose a new biological term to encapsulate this idea: evo-engineering. We believe that NMps offer an ideal experimental system in which to approach this concept.

In order to test the validity of this approach and to probe the key parameters that are regulated to generate the observed differences between model organisms, we require access to dynamic information at the single cell level. Given their transparency and ease of accessibility, this is possible in zebrafish, which also have the advantage of being able to explore the molecular mechanisms leading to NMp bipotency and cell fate decisions in the absence of overt self-renewal. This is highly complementary to the study of the mouse embryo for which we have good fate mapping and retrospective clonal analysis data (Tzouanacou et al., 2009; Wymeersch et al., 2016). However, live imaging is still problematic for post-implantation stages in the mouse, particularly for non-superficial tissue level events.

An interesting recent development for our understanding of the molecular mechanisms establishing and controlling the behaviour of the NMps is the emergence of Embryonic Stem Cells (ESCs) as an experimental system to study cell fate decision and the emergence of tissues and organs in mammalian development. Calibration of differentiation conditions of mouse ESCs has allowed the generation, in adherent culture, of cells with gene expression profiles similar to NMps (Gouti et al., 2014; Tsakiridis et al., 2014; Turner et al., 2014). These NMp-like (NMpl) cells coexpress T/Bra and Sox2, exhibit heterogeneities in the expression of a variety of other genes associated with the progenitors and, as a population, are able to differentiate into neural and paraxial mesoderm in a Wnt and FGF dependent manner. A similar population has been obtained from human ES cells with slightly different culture conditions (Gouti et al., 2015; Lippmann et al., 2015). In one case (Lippmann et al., 2015), the population has been maintained for a few days. However, it is not yet clear to what degree these cells reflect the NMps in the embryo as despite the similarities with their in vivo counterparts, they are difficult to maintain in culture and clonal analysis has revealed that very few of them have the property of self-renewal and differentiation. Furthermore, in the adherent culture Wnt signalling favours mesodermal differentiation, something that has been observed in vivo, but suppresses neural fates, something which has not observed in the embryo (Garriock et al., 2015; Wymeersch et al., 2016). A complementary alternative to the adherent culture is provided by a novel 3D non adherent culture system in which cells form embryo like structure and develop an NMp population that can promote organized elongation (Turner et al., 2016; van den Brink et al., 2014). The combination of adherent culture and this system will lead to understanding of the NMps.

The study of NMps is important not only because of their central role in generating the embryonic body axis, but also because of the potential generation of spinal cord neurons in culture for the study of as-yet difficult to model diseases such as spinal cord atrophy. Key to achieving this goal is the development of reliable clonal expansion and differentiation strategies for the generation of spinal cord precursors *in vivo*. Given the differences in cellular mechanisms that generate the spinal cord in amniotes is largely due to an increased in the clonal expansion of NMps, a continued comparative analysis of these processes as outlined above may well bring some practical insight into how to develop such strategies. Furthermore, the comparative analysis of NMp developmental dynamics represents an ideal experimental system in which to approach a deep and elusive problem in evolutionary developmental biology: that of how to generate diverse behaviours at the cellular level from seemingly conserved regulatory processes at the level of the whole population.

## Acknowledgements

The research of BS was funded by a Marie Curie fellowship

